# Hippocampal inputs engage CCK+ interneurons to mediate endocannabinoid-modulated feed-forward inhibition in the prefrontal cortex

**DOI:** 10.1101/2020.03.12.988881

**Authors:** Xingchen Liu, Jordane Dimidschstein, Gordon Fishell, Adam G. Carter

## Abstract

Connections from the ventral hippocampus (vHPC) to the prefrontal cortex (PFC) regulate cognition, emotion and memory. These functions are also tightly regulated by inhibitory networks in the PFC, whose disruption is thought to contribute to mental health disorders. However, relatively little is known about how the vHPC engages different populations of interneurons in the PFC. Here we use slice physiology and optogenetics to study vHPC-evoked feed-forward inhibition in the mouse PFC. We first show that cholecystokinin (CCK+), parvalbumin (PV+), and somatostatin (SOM+) interneurons are prominent in layer 5 (L5) of infralimbic PFC. We then show that vHPC inputs primarily activate CCK+ and PV+ interneurons, with weaker connections onto SOM+ interneurons. CCK+ interneurons make stronger synapses onto pyramidal tract (PT) cells over nearby intratelencephalic (IT) cells. However, CCK+ inputs undergo depolarization-induced suppression of inhibition (DSI) and CB1 receptor modulation only at IT cells. Moreover, vHPC-evoked feed-forward inhibition undergoes DSI only at IT cells, confirming a central role for CCK+ interneurons. Together, our findings show how vHPC directly engages multiple populations of inhibitory cells in deep layers of the infralimbic PFC, highlighting unexpected roles for both CCK+ interneurons and endocannabinoid modulation in hippocampal-prefrontal communication.

## INTRODUCTION

The prefrontal cortex (PFC) controls cognitive and emotional behaviors (Euston et al., 2012; Miller and Cohen, 2001) and is disrupted in many neuropsychiatric disorders (Godsil et al., 2013; Sigurdsson and Duvarci, 2015). PFC activity is driven and maintained by long-range glutamatergic inputs from a variety of other brain regions (Hoover and Vertes, 2007; Miller and Cohen, 2001). Strong, unidirectional connections from the ventral hippocampus (vHPC) contribute to both working memory and threat conditioning in rodents (Jones and Wilson, 2005; Sotres-Bayon et al., 2012; Spellman et al., 2015). Dysfunction of vHPC to PFC connectivity is also implicated in schizophrenia, anxiety disorders, chronic stress disorders, and depression (Godsil et al., 2013; Sigurdsson and Duvarci, 2015). To understand these roles, it is necessary to establish how vHPC inputs engage local excitatory and inhibitory networks within the PFC.

The vHPC primarily projects to the ventral medial PFC in rodents, with axons most prominent in layer 5 (L5) of infralimbic (IL) PFC (Phillips et al., 2019). These excitatory inputs contact multiple populations of pyramidal cells, and are much stronger at intratelencephalic (IT) cells than nearby pyramidal tract (PT) cells (Liu and Carter, 2018). vHPC inputs can drive robust firing of IT cells (Liu and Carter, 2018), which may be important for maintaining activity during behavioral tasks (Padilla-Coreano et al., 2016; Spellman et al., 2015). However, they also evoke prominent feed-forward inhibition at pyramidal cells (Marek et al., 2018), with excitation and inhibition evolving with different dynamics (Liu and Carter, 2018). Here we focus on the mechanisms responsible for this inhibition by establishing which interneurons are engaged by vHPC inputs to the PFC.

As in other cortices, PFC activity is regulated by a variety of GABAergic interneurons, which have distinct functions (Abbas et al., 2018; Courtin et al., 2014; Tremblay et al., 2016). Parvalbumin-expressing (PV+) interneurons mediate feed-forward inhibition via strong synapses at the soma of pyramidal cells (Atallah et al., 2012; Cruikshank et al., 2007; Gabernet et al., 2005). In contrast, somatostatin-expressing (SOM+) interneurons mediate feed-back inhibition via facilitating synapses onto the dendrites (Gentet et al., 2012; Silberberg and Markram, 2007). However, during trains of repetitive activity, SOM+ interneurons can also participate in feed-forward inhibition (McGarry and Carter, 2016; Tan et al., 2008). Interestingly, recent *in vivo* studies suggest both PV+ and SOM+ interneurons may contribute to vHPC-evoked inhibition in the PFC (Abbas et al., 2018; Marek et al., 2018).

While PV+ and SOM+ interneurons are prominent in the PFC, there is also an unusually high density of cholecystokinin-expressing (CCK+) interneurons (Whissell et al., 2015). In the hippocampus, these soma-targeting inhibitory cells help tune the motivation and emotional state of animals (Armstrong and Soltesz, 2012; Freund, 2003; Freund and Katona, 2007). They also highly express cannabinoid type 1 (CB1) receptors on their axon terminals (Katona et al., 1999), and can be strongly modulated by endocannabinoids (Wilson et al., 2001; Wilson and Nicoll, 2001). For example, brief depolarization of postsynaptic pyramidal cells releases endocannabinoids that bind to CB1 receptors on CCK+ axon terminals and inhibit presynaptic GABA release, a process known as depolarization-induced suppression of inhibition (DSI) (Wilson and Nicoll, 2001).

While CCK+ interneurons are prominent in the PFC and may play a role in different forms of inhibition, they remain relatively understudied. A major technical reason is low-level expression of CCK in pyramidal cells (Taniguchi et al., 2011), which makes CCK+ interneurons challenging to specifically target. For example, expressing Cre-dependent reporters in CCK-Cre transgenics also label pyramidal cells in the cortex (Taniguchi et al., 2011). Fortunately, this challenge can be overcome with intersectional viruses using the Dlx enhancer, which restricts expression to interneurons (Dimidschstein et al., 2016). This approach allows identification of CCK+ interneurons, enabling targeted recordings and optogenetic access to study their connectivity and modulation in the PFC.

Here we examine vHPC-evoked inhibition at L5 pyramidal cells in IL PFC using slice physiology, optogenetics and intersectional viral tools. We find vHPC inputs activate PV+, SOM+, and CCK+ interneurons, with different dynamics during repetitive activity. Inputs to PV+ and CCK+ interneurons are strong but depressing, while those onto SOM+ interneurons are weak but facilitating. CCK+ interneurons contact L5 pyramidal cells, with stronger connections onto PT cells than neighboring IT cells. However, endocannabinoid modulation via DSI and direct activation of CB1 receptors only occurs at synapses onto IT cells. Endocannabinoid modulation of vHPC-evoked feed-forward inhibition also occurs only at IT cells, highlighting a central role for CCK+ interneurons. Together, our findings show how the vHPC engages interneurons to inhibit the PFC, while revealing a novel property of cell-type specific endocannabinoid modulation in this circuit.

## MATERIALS AND METHODS

### Animals

Experiments used wild-type and transgenic mice of either sex in a C57BL/6J background (all breeders from Jackson Labs). Homozygote male breeders (PV-Cre = JAX 008069, SOM-Cre = JAX 013044, CCK-Cre = JAX 012706) were paired with female wild-type or Ai14 breeders (JAX 007914) to yield heterozygote offspring for experiments. All experimental procedures were approved by the University Animal Welfare Committee of New York University.

### Viruses

AAV viruses used in this study were: AAV1.EF1a.DIO.hChR2(H134R)-eYFP.WPRE.hGH (UPenn AV-1-20298P), AAV1.hysn.hChR2(H134R)-eYFP.WPRE.hGH (UPenn AV-26973P), AAV1.EF1a.DIO.eYFP.WPRE.hGH (Upenn AV-1-27056), AAVrg.CAG.GFP (Addgene 37825-AAVrg), AAVrg.CAG.tdTomato (Addgene 59462-AAVrg). Additional viral constructs were assembled for Cre-dependent expression of a reporter under the control of the Dlx5/6 enhancer: AAV-Dlx-Flex-GFP (Addgene #83900) and AAV-Dlx-Flex-ChR2-mCherry (Dimidschstein et al., 2016). These constructs take advantage of the double-floxed inverted system, in which two consecutive and incompatible lox-sites are placed both in 5’ and 3’ of the reversed coding sequences of the viral reporter, restricting expression to interneurons.

### Stereotaxic injections

Mice aged 4-6 weeks were deeply anesthetized with either isoflurane or a mixture of ketamine and xylazine, then head-fixed in a stereotaxic (Kopf Instruments). A small craniotomy was made over the injection site, using these coordinates relative to Bregma: PFC = ±0.4, −2.3, +2.1 mm, PAG = −0.6, both −2.5 and −3, −4.0 mm, vHPC = −3.3, both −3.6 and −4.2, −3 mm (mediolateral, dorsoventral, and rostrocaudal axes). For retrograde labeling, pipettes were filled with red retrogradely transported fluorescent beads (Lumafluor), Cholera Toxin Subunit B (CTB) conjugated to Alexa 647 (Life Technologies), or viruses. Borosilicate pipettes with 5 to 10 µm diameter tips were back-filled with dye and/or virus, and a volume of 130-550 nl was pressure-injected using a Nanoject III (Drummond) every 30 s. The pipette was left in place for an additional 5 min, allowing time to diffuse away from the pipette tip, before being slowly retracted from the brain. For both retrograde and viral labeling, animals were housed for 2-3 weeks before slicing.

### Histology and fluorescence microscopy

Mice were anesthetized with a lethal dose of ketamine and xylazine, then perfused intracardially with 0.01 M phosphate buffered saline (PBS) followed by 4% paraformaldehyde (PFA) in 0.01 M PBS. Brains were fixed in 4% PFA in 0.01 M PBS overnight at 4°C. Slices were prepared at a thickness of 70 μm for imaging intrinsic fluorescence or 40 µm for antibody staining (Leica VT 1000S vibratome). For immunohistochemistry, slices were incubated with blocking solution (1% bovine serum albumin and 0.2% Triton-X in 0.01 m PBS) for 1 hour at room temperature before primary antibodies were applied in blocking solution [mouse anti-parvalbumin antibody (Millipore, MAB1572) at 1:2000 overnight, rat anti-somatostatin (Millipore, MAB354) at 1:200 overnight, guinea pig anti-CB1R (Frontier Institute, Af530) at 1:500 for 36 hours] at 4 °C. Slices were then incubated with secondary antibodies in blocking solution [goat anti-mouse 647 at 1:200, goat anti-rat 647 at 1:200, goat anti-guinea pig 647 at 1:500 (Invitrogen)] for 1.5 h at room temperature before mounted under glass coverslips on gelatin-coated slides using ProLong Gold antifade reagent with DAPI (Invitrogen). Images were acquired using a confocal microscope (Leica SP8). Image processing involved adjusting brightness, contrast, and manual cell counting using ImageJ (NIH).

### Slice preparation

Mice aged 6-8 weeks were anesthetized with a lethal dose of ketamine and xylazine, and perfused intracardially with ice-cold external solution containing the following (in mM): 65 sucrose, 76 NaCl, 25 NaHCO_3_, 1.4 NaH_2_PO_4_, 25 glucose, 2.5 KCl, 7 MgCl_2_, 0.4 Na-ascorbate, and 2 Na-pyruvate (295–305 mOsm), and bubbled with 95% O_2_/5% CO_2_. Coronal slices (300 μm thick) were cut on a VS1200 vibratome (Leica) in ice-cold external solution, before being transferred to ACSF containing (in mM): 120 NaCl, 25 NaHCO_3_, 1.4 NaH_2_PO_4_, 21 glucose, 2.5 KCl, 2 CaCl_2_, 1 MgCl_2_, 0.4 Na-ascorbate, and 2 Na-pyruvate (295–305 mOsm), bubbled with 95% O_2_/5% CO_2_. Slices were kept for 30 min at 35°C, before being allowed to recover for 30 min at room temperature before starting recordings. All recordings were conducted at 30−32°C.

### Electrophysiology

Whole-cell recordings were obtained from pyramidal neurons or interneurons located in layer 5 (L5) of infralimbic (IL) PFC. Neurons were identified by infrared-differential interference contrast or fluorescence, as previously described (Chalifoux and Carter, 2010). In the case of pyramidal cells, projection target was established by the presence of retrobeads or Alexa-conjugated CTB, as previously described (Little and Carter, 2013). Pairs of adjacent cells were chosen for sequential recording, ensuring they received similar inputs (typically < 50 µm between cells). Borosilicate pipettes (2–5 MΩ) were filled with internal solutions. Three types of recording internal solutions were used. For current-clamp recordings (in mM): 135 K-gluconate, 7 KCl, 10 HEPES, 10 Na-phosphocreatine, 4 Mg_2_-ATP, and 0.4 Na-GTP, 290–295 mOsm, pH 7.3, with KOH. For voltage-clamp recordings (in mM): 135 Cs-gluconate, 10 HEPES, 10 Na-phosphocreatine, 4 Mg_2_-ATP, and 0.4 Na-GTP, 0.5 EGTA, 10 TEA-chloride, and 2 QX314, 290–295 mOsm, pH 7.3, with CsOH. For DSI experiments (in mM): 130 K-gluconate, 1.5 MgCl_2_, 10 HEPES, 1.1 EGTA, 10 phosphocreatine, 2 MgATP, 0.4 NaGTP. In some experiments studying cellular morphology, 5% biocytin was also included in the recording internal solution. After allowing biocytin to diffuse through the recorded cell for at least 30 minutes, slices were fixed with 4% PFA before staining with streptavidin conjugated to Alexa 647 (Invitrogen).

Electrophysiology recordings were made with a Multiclamp 700B amplifier (Axon Instruments), filtered at 4 kHz for current-clamp and 2 kHz for voltage-clamp, and sampled at 10 kHz. The initial series resistance was <20 MΩ, and recordings were ended if series resistance rose above 25 MΩ. In some experiments, 1 μM TTX was added to block APs, along with 100 μM 4-AP and 4 mM external Ca^2+^ to restore presynaptic release. In many experiments, 10 μM CPP was used to block NMDA receptors. In current-clamp experiments characterizing intrinsic properties, 10 μM NBQX, 10 μM CPP and 10 µM gabazine were used to block excitation and inhibition. In some experiments, 10 µm AM-251 was used to block CB1 receptors or 1 µM WIN 55,212-2 was used to activate CB1 receptors. All chemicals were purchased from either Sigma or Tocris Bioscience.

### Optogenetics

Channelrhodopsin-2 (ChR2) was expressed in presynaptic neurons and activated with a brief light pulse from a blue LED (473 nm) (Thorlabs). For wide-field illumination, light was delivered via a 10x 0.3NA objective (Olympus) centered on the recorded cell. LED power was routinely calibrated at the back aperture of the objective. LED power and duration were adjusted to obtain reliable responses, with typical values of 0.4 to 10 mW and 2 ms, respectively.

### Data analysis

Electrophysiology and imaging data were acquired using National Instruments boards and custom software in MATLAB (MathWorks). Off-line analysis was performed using Igor Pro (WaveMetrics). Intrinsic properties were determined as follows. Input resistance was calculated from the steady-state voltage during a −50 pA, 500 ms current step. Voltage sag ratio was calculated as (V_sag_ − V_ss_) / (V_sag_ − V_baseline_), where V_sag_ is average over a 1 ms window around the minimum, V_ss_ is average of last 50 ms, and V_baseline_ is average of 50 ms preceding the current injection. The membrane time constant (tau) was measured using exponential fits to these hyperpolarizations. Adaptation was calculated as the ratio of the first and last inter-spike intervals, such that a value of 1 indicates no adaptation and values <1 indicate lengthening of the inter-spike interval. For experiments with a single optogenetic stimulation, the PSC amplitude was measured as the average value across 1 ms around the peak subtracted by the average 100 ms baseline value prior to the stimulation. For experiments with a train of optogenetic stimulation, each PSC amplitude was measured as the average value in a 1 ms window around the peak, minus the average 2 ms baseline value before each stimulation. Most summary data are reported in the text and figures as arithmetic mean ± SEM. Ratios of responses at pairs of cells are reported as geometric mean in the text, and with ± 95% confidence interval (CI) in the figures, unless otherwise noted. Comparisons between unpaired data were performed using non-parametric Mann-Whitney test. Comparisons between data recorded in pairs were performed using non-parametric Wilcoxon test. Two-tailed p values <0.05 were considered significant.

## RESULTS

### vHPC-evoked inhibition and interneurons in L5 of infralimbic PFC

We studied vHPC-evoked inhibition at pyramidal neurons using whole-cell recordings and optogenetics in acute slices of the mouse PFC. To allow for visualization and activation of inputs to the PFC, we injected ChR2-expressing virus (AAV-ChR2-EYFP) into the ipsilateral vHPC (**Fig. 1A**) (Little and Carter, 2012). In the same animals, we labeled IT cells by co-injecting retrogradely transported, fluorescently-tagged cholera toxin subunit B (CTB) into the contralateral PFC (cPFC) (**Fig. 1A**) (Anastasiades et al., 2018). After 2-3 weeks of expression, we prepared *ex vivo* slices of the medial PFC, observing vHPC axons and IT cells in L5 of IL PFC (**Fig. 1B**) (Liu and Carter, 2018). We then recorded in voltage-clamp from IT cells and activated vHPC inputs using wide-field illumination (5 pulses at 20 Hz, 2 ms pulse duration). Trains of vHPC inputs evoked excitatory postsynaptic currents (EPSCs) at −65 mV and inhibitory postsynaptic currents (IPSCs) at +15 mV (**Fig. 1C**; EPSC_1_ = 386 **±** 49 pA, IPSC_1_ = 961 **±** 207 pA, E/I = 0.50 ±0.08; n = 7 cells, 3 animals). These findings indicate that vHPC inputs drive robust feed-forward inhibition in deep layers of the IL PFC, motivating us to identify which interneurons are responsible.

**Figure 1:**
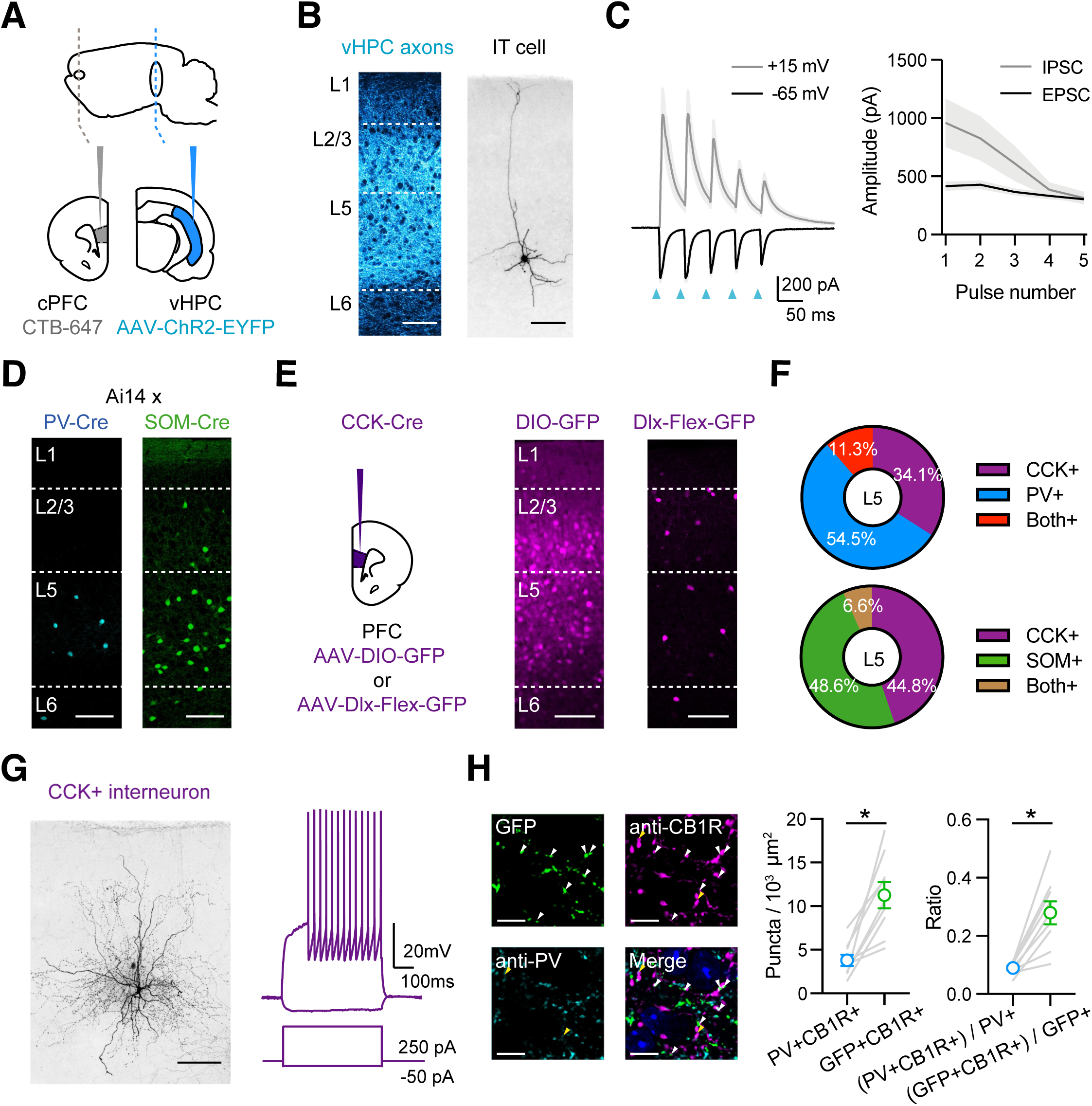
vHPC-evoked feed-forward inhibition and CCK+ interneurons. **A)** Schematic for injections of AAV-ChR2-EYFP into vHPC and CTB-647 into cPFC. **B)** *Left,* Confocal image of vHPC axons (blue) in IL PFC. Scale bar = 100 µm. *Right,* Confocal image of biocytin-filled L5 IT cell in IL PFC. Scale bar = 100 µm. **C)** *Left,* Average vHPC-evoked EPSCs at −65 mV (black) and IPSCs at +15 mV (grey). Blue arrows = 5 pulses at 20 Hz. *Right,* Average response amplitudes as function of pulse number (n = 7 cells, 3 animals). **D)** Td-tomato labeling of PV+ (blue) and SOM+ (green) interneurons in PV-Cre x Ai14 and SOM-Cre x Ai14 animals, respectively. **E)** *Left*, Schematic for injections of viruses into PFC of CCK-Cre mouse. *Middle*, Injection of AAV-DIO-GFP labels CCK+ interneurons and pyramidal cells (n = 3 animals). *Right*, Injection of AAV-Dlx-Flex-GFP labels CCK+ interneurons (n = 3 animals). Scale bars = 100 µm. **F)** Quantification of co-labeling of CCK+ interneurons with PV (top) and SOM (bottom) (PV staining, n = 308 cells, 17 slices, 6 animals; SOM staining, n = 105 cells, 8 slices, 3 animals). **G)** *Left*, Confocal image of a biocytin-filled CCK+ interneuron in L5 of IL PFC. Scale bar = 100 µm. *Right*, Response to positive and negative current injections. **H)** *Left,* Confocal images of GFP (green), anti-CB1R staining (purple), anti-PV staining (blue), and merge. Arrow heads: white = GFP+CB1R+ co-labeling, yellow = PV+CB1R+ co-labeling. *Middle,* Quantification of PV+CB1R+ and GFP+CB1R+ quanta per 10^3^ µm^2^ (n = 9 sections, 3 animals). *Right,* Quantification of the ratios of CB1R+ puncta among PV+ and GFP+ puncta. Each line represents counts from one slice.

In principle, feed-forward inhibition could be mediated by a variety of interneurons, including PV+ and SOM+ interneurons (Abbas et al., 2018; Anastasiades et al., 2018; Marek et al., 2018; McGarry and Carter, 2016). To visualize these cells in the PFC, we crossed PV-Cre and SOM-Cre mice with reporter mice (Ai14) that express Cre-dependent tdTomato (Hippenmeyer et al., 2005; Madisen et al., 2010; Taniguchi et al., 2011). We observed labeling of PV+ and SOM+ interneurons in L5 of IL PFC, suggesting they could be contacted by vHPC afferents (**Fig. 1D**). However, the PFC also has a high density of cholecystokinin-expressing (CCK+) interneurons (Whissell et al., 2015), which mediate inhibition in other cortices and the hippocampus (Armstrong and Soltesz, 2012; Freund and Katona, 2007), and could also participate in the PFC. To label these cells, we initially injected AAV-DIO-GFP into CCK-Cre mice but observed labeling of both interneurons and pyramidal neurons across multiple layers (**Fig. 1E**). To restrict labeling to CCK+ interneurons, we instead used AAV-Dlx-Flex-GFP, expressing Cre-dependent GFP under control of the Dlx enhancer (Dimidschstein et al., 2016). Injecting AAV-Dlx-Flex-GFP selectively labeled CCK+ interneurons in the IL PFC, including prominent labeling in L5 (**Fig. 1E**). Importantly, we found little co-labeling of CCK+ cells with either PV (**Fig. 1F & S1A;** 11.3 % overlap, n = 308 cells total, 17 slices, 6 animals) or SOM (6.6% overlap, n = 105 cells total, 8 slices, 3 animals).

To confirm targeting of CCK+ interneurons, we next used whole-cell recordings followed by post-hoc reconstructions (**Fig. 1G**). We found both axons and dendrites in L5 of IL PFC, and intrinsic properties similar to reports in other parts of the brain (**Fig. S1B;** Rin = 130 ± 13 MΩ, Vm = −68 ± 1 mV, Sag = 4.1 ± 1.0 %, Adaptation = 0.81 ± 0.04, Tau = 9.1 ± 0.6 ms; n = 12 cells, 4 animals) (Daw et al., 2009). CCK+ interneurons are also distinguished from PV+ cells by enrichment of the cannabinoid type 1 receptor (CB1R) on presynaptic terminals (Bodor et al., 2005; Dudok et al., 2015; Katona et al., 1999). To examine expression in the IL PFC, we used immunocytochemistry and detected CB1R+ puncta located at either PV+ or CCK+ axons (**Fig. 1H**). We found CB1R+ puncta showed higher co-localization with CCK+ axons (based on GFP+ expression) than with PV+ axons (**Fig. 1H**, puncta density in 10^3^/µm^2^: PV+CB1R+ = 3.79 ± 0.64, GFP+CB1R+ = 11.27 ± 1.5, p = 0.0003). These results confirm our viral strategy can identify CCK+ interneurons, which are present in deep layers of IL. Moreover, they suggest PV+, SOM+, and CCK+ interneurons are positioned to receive vHPC inputs and may mediate feed-forward inhibition.

### vHPC inputs primarily engage PV+ and CCK+ interneurons

Previous studies have suggested that vHPC inputs engage PV+ and SOM+ interneurons in the PFC (Abbas et al., 2018; Marek et al., 2018). However, these connections have not been examined in L5 of IL, where vHPC connections are strongest (Liu and Carter, 2018). Moreover, little is known about the activation of CCK+ interneurons, which may play distinct functional roles (Armstrong and Soltesz, 2012; Freund and Katona, 2007). To study connectivity, we injected AAV-ChR2-YFP into the vHPC and recorded EPSCs from identified interneurons. To compare across animals and cell types, we recorded from neighboring IT cells within the same slice, which receive the bulk of vHPC inputs (Liu and Carter, 2018). To isolate monosynaptic connections, we included TTX (1 μM), 4-AP (10 μM), and elevated Ca^2+^ (4 mM), which blocks action potentials (APs) but restores presynaptic release (Little and Carter, 2012; Petreanu et al., 2009). We also kept the light intensity and duration constant within pairs, allowing us to account for differences in viral expression across animals and slices (Liu and Carter, 2018). We found that vHPC inputs evoked robust EPSCs at CCK+ interneurons, which were similar in amplitude to IT cells (**Fig. 2A**; IT = 674 **±** 102 pA, CCK+ = 413 **±** 89 pA, CCK+/IT ratio = 0.56, p = 0.13; n = 9 pairs, 4 animals). We also found prominent vHPC-evoked EPSCs at PV+ interneurons, with comparable amplitudes at IT cells (**Fig. 2B**; IT = 569 **±** 91 pA, PV+ = 555 **±** 83 pA, PV+/IT ratio = 0.94, p = 0.82; n = 9 pairs, 4 animals). In contrast, vHPC inputs evoked much smaller EPSCs at SOM+ interneurons compared to IT cells (**Fig. 2C**; IT = 421 **±** 69 pA, SOM+ = 95 **±** 33 pA, SOM+/IT ratio = 0.17, p = 0.004; n = 9 pairs, 4 animals). These findings show that all three cell types receive direct vHPC inputs, with greater responses at CCK+ and PV+ interneurons.

**Figure 2:**
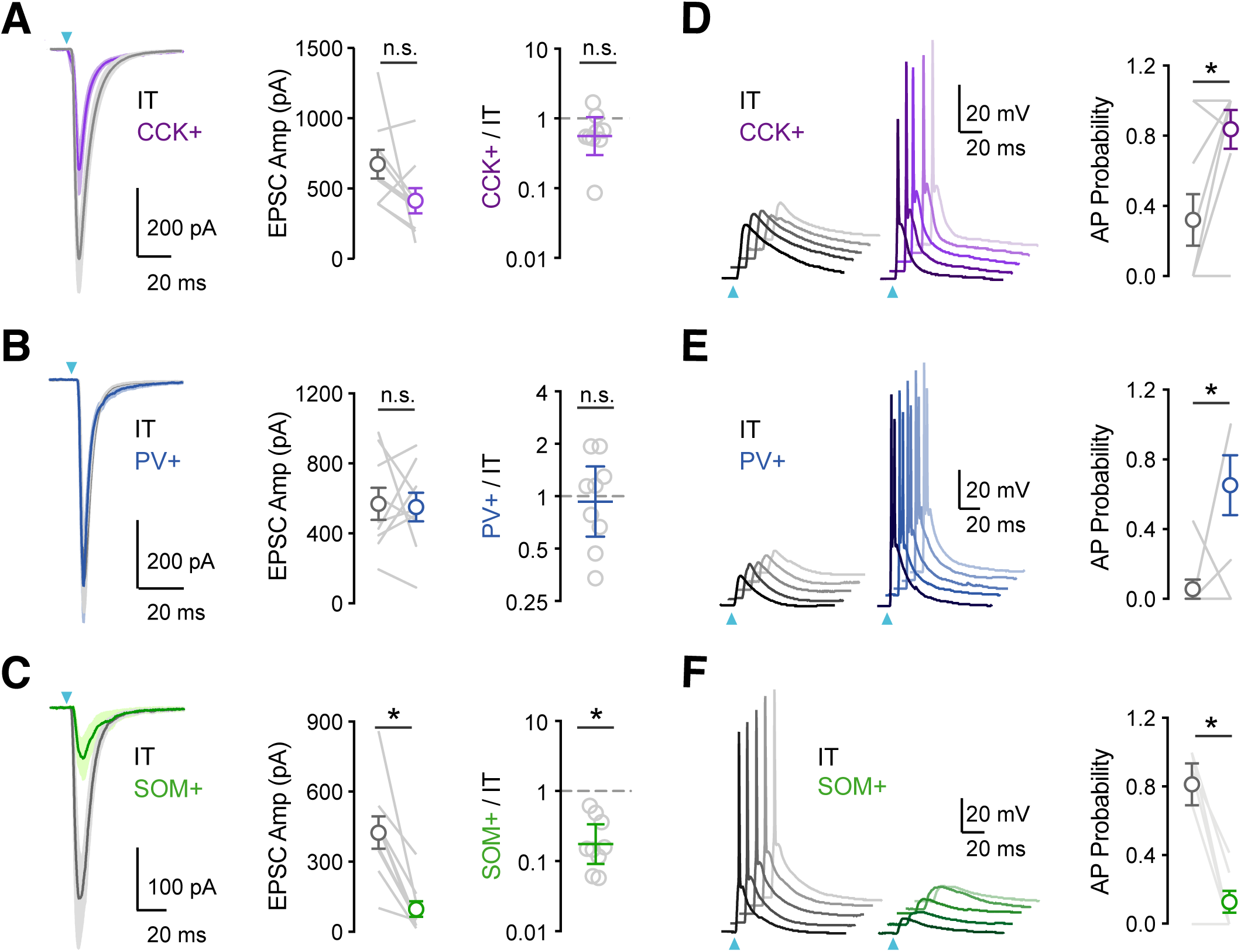
vHPC inputs differentially engage PV+, SOM+ and CCK+ interneurons. **A)** *Left,* Average vHPC-evoked EPSCs at pairs of L5 IT (grey) and CCK+ (purple) cells in IL PFC. Blue arrow = light pulse. *Middle*, Summary of EPSC amplitudes. *Right*, Summary of CCK+ / IT EPSC amplitude ratios (n = 9 pairs, 4 animals). **(B – C)** Similar to (A) for pairs of IT and PV+ cells (n = 9 pairs, 4 animals) or pairs of IT and SOM+ cells (n = 9 pairs, 4 animals). **D)** *Left*, vHPC-evoked EPSPs and APs recorded in current-clamp from resting membrane potential at pairs of L5 IT (grey) and CCK+ (purple) cells in IL PFC, with 5 traces offset for each cell. *Right*, Summary of AP probability at pairs of IT and CCK+ cells. Blue arrow = 3.5 mW light pulse (n = 9 pairs, 4 animals). **(E – F)** Similar to (D) for pairs of IT and PV+ cells (3.5 mW light pulses, n = 8 pairs, 4 animals) or pairs of IT and SOM+ cells (4.8 mW light pulses, n = 7 pairs, 3 animals). * p<0.05

Having established the targeting of vHPC inputs, we next accessed their ability to drive action potentials (APs) at the three classes of interneurons. We used similar viral and labeling approaches, but in this case conducted current-clamp recordings in the absence of TTX and 4-AP. We also kept light intensity constant within each set of experiments, making sure we evoked action potentials in at least one of the recorded pair of cells. We found that single vHPC inputs evoked APs in pairs of CCK+ and IT cells, but the interneurons showed higher probability of firing (**Fig. 2D**; AP probability: IT = 0.32 ± 0.15, CCK+ = 0.84 ± 0.11, p = 0.03; n = 9 pairs, 4 animals). Similarly, we observed that vHPC inputs also preferentially activate PV+ over IT cells (**Fig. 2E**; AP probability: IT = 0.06 ± 0.06, PV+ = 0.65 ± 0.17, p = 0.04; n = 8 pairs, 4 animals). In contrast, SOM+ interneurons remained unresponsive to vHPC inputs, even when using higher light intensities that were able to activate IT cells (**Fig. 2F**; AP probability: IT = 0.81 ± 0.12, SOM+ = 0.13 ± 0.06, p = 0.01; n = 7 pairs, 3 animals). These results established a hierarchy for activation, suggesting CCK+ interneurons are engaged by vHPC inputs and mediate feed-forward inhibition.

### Short-term dynamics differ between populations of interneurons

Repetitive activity at vHPC to PFC connections is functionally important and depends on stimulus frequency (Liu and Carter, 2018; Siapas et al., 2005). At the synaptic level, repetitive activity engages short-term plasticity to change the strength of individual connections (Zucker and Regehr, 2002). We next examined the response to repetitive vHPC inputs by stimulating with brief trains (5 pulses at 20Hz) in the absence of TTX and 4-AP (Liu and Carter, 2018; McGarry and Carter, 2016). We observed that EPSCs at CCK+ and PV+ interneurons strongly depress over the course of stimulus trains (**Fig. 3A, B & D**; EPSC_2_ / EPSC_1_: CCK+ = 0.82 ± 0.06; n = 6 cells, 3 animals; PV+ = 0.88 **±** 0.06; n = 7 cells, 3 animals). In contrast, the EPSCs at SOM+ interneurons initially facilitated during trains (**Fig. 3C & D**; EPSC_2_ / EPSC_1_ = 1.78 ± 0.23; n = 8 cells, 3 animals). These results indicate the vHPC engages all three cell types, with connections at PV+ and CCK+ interneurons strong but depressing, and those at SOM+ interneurons weak but facilitating.

**Figure 3:**
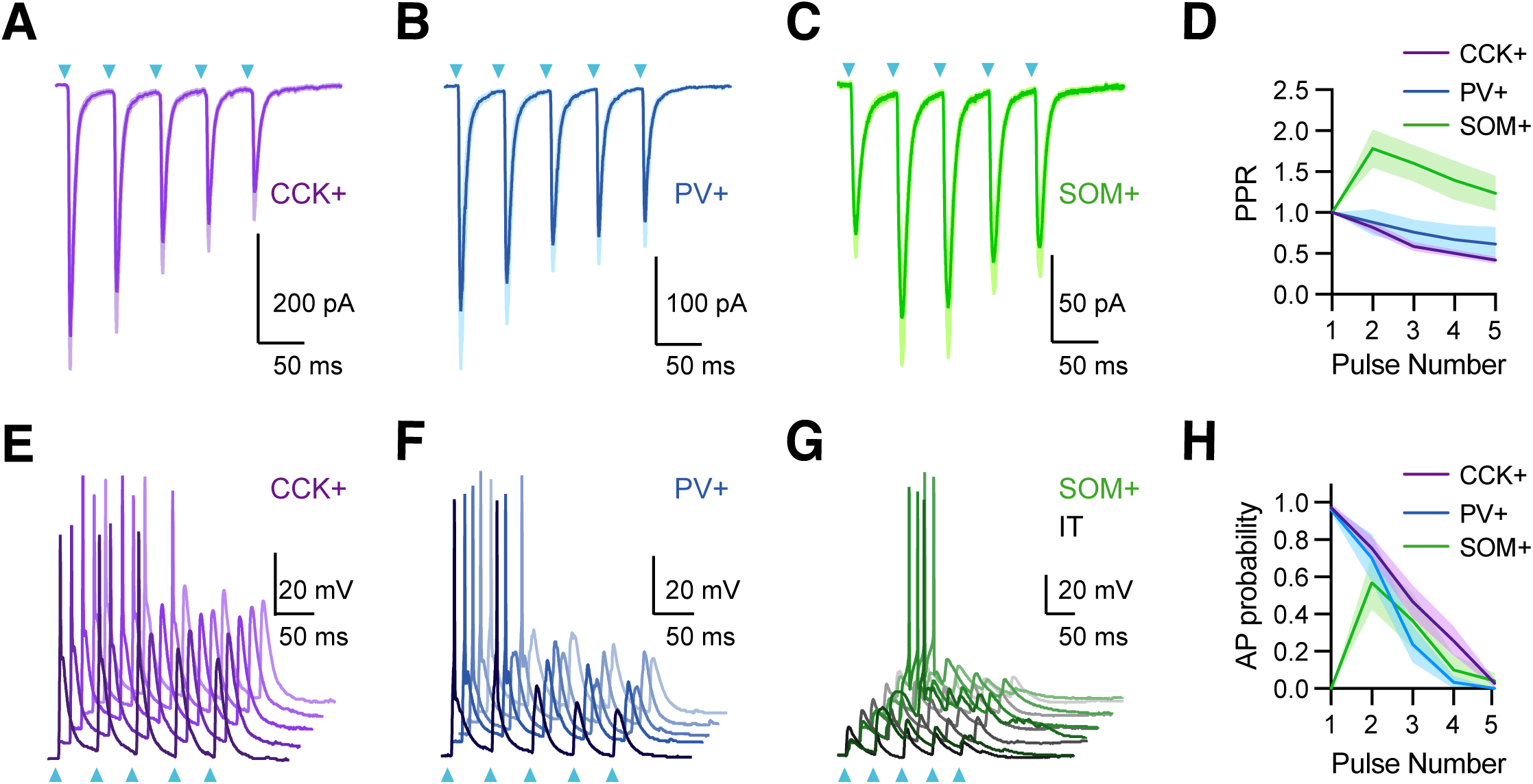
vHPC inputs drive interneurons with distinct temporal dynamics. **A)** Average vHPC-evoked EPSCs at CCK+ interneurons, recorded in voltage-clamp at −65 mV (5 pulses at 20 Hz). Blue arrows = light pulses (n = 6 cells, 3 animals). **B)** Similar to (A) for PV+ interneurons (n = 7 cells, 3 animals). **C)** Similar to (A) for SOM+ interneurons (n = 8 cells, 3 animals). **D)** Average paired-pulse ratio (PPR) of vHPC-evoked EPSCs at CCK+, PV+, and SOM+ interneurons. **E)** vHPC-evoked EPSPs and APs, recorded in current-clamp from resting membrane potential at an example CCK+ interneuron in L5 of IL PFC, with 5 traces offset for the cell. Blue arrows = light pulses (n = 12 cells, 5 animals). **F)** Similar to (E) for PV+ interneurons (n = 6 cells, 3 animals). **G)** Similar to (E) for SOM+ interneurons. Note that each SOM+ cell was studied with a nearby IT cell under same recording and stimulation conditions, where the absence of AP firing at IT cells indicated subthreshold of network recurrent activation (n = 6 cells, 3 animals). **H)** Summary of average AP probability as a function of pulse number for CCK+, PV+, and SOM+ interneurons. * p<0.05

The short-term dynamics of vHPC inputs suggested differential engagement of PV+, SOM+, and CCK+ interneurons during repetitive activity. We observed that stimulus trains of vHPC inputs generated decreasing AP probabilities at CCK+ and PV+ interneurons (**Fig. 3E, F & H**; stimulus 1 & 2 AP probabilities: CCK+ = 0.97 ± 0.02 & 0.75 ± 0.06; n = 12 cells, 5 animals; PV+ = 0.96 ± 0.02 & 0.70 ± 0.13; n = 6 cells, 3 animals;). In contrast, although SOM+ interneurons did not fire with single vHPC inputs, they were activated during trains. Importantly, these responses were not due to recurrent network activity, as stimulation intensity was chosen here to ensure that IT cells remained quiescent (**Fig. 3G & H;** stimulus 1 & 2 AP probabilities: IT = 0 & 0, SOM+ = 0 & 0.57 ± 0.14; n = 6 pairs, 4 animals). These findings suggest the vHPC differentially engages three interneurons with distinct dynamics during repetitive activity, with strong and depressing PV+ and CCK+ activity contributing early, and weaker but facilitating SOM+ activity contributing later.

### CCK+ interneurons make connections onto L5 pyramidal cells

Our results indicate that vHPC inputs strongly engage CCK+ interneurons in the PFC, suggesting a role in feed-forward inhibition. However, the connections made by CCK+ interneurons onto nearby pyramidal cells are not well established in the cortex. To study CCK+ output, we developed a new virus (AAV-Dlx-Flex-ChR2-mCherry) to express ChR2 in a Cre-dependent manner under the Dlx enhancer (see Methods). We injected this virus into the PFC of CCK-Cre mice to selectively express ChR2 in CCK+ interneurons (**Fig. 4A**). Whole-cell current-clamp recordings showed labeled cells could be rapidly and reliably activated by brief pulses of blue light (**Fig. 4A**). To study inhibitory connections, we then recorded CCK+-evoked currents from unlabeled L5 pyramidal cells in IL PFC (**Fig. 4B**). To detect both EPSCs or IPSCs, we used a low chloride internal and held at −50 mV, such that inward currents were EPSCs and outward currents were IPSCs (Glickfeld and Scanziani, 2006). We observed robust CCK+-evoked IPSCs, which were unaffected by blockers of AMPAR (10 µM NBQX) and NMDAR (10 µM CPP) but abolished by blockers of GABA_A_R (10 µM gabazine) (**Fig. 4B**; ACSF = 97 ± 21 pA, NBQX+CPP = 97 ± 21 pA, gabazine = 0.4 ± 0.2 pA; ACSF vs NBQX + CPP, p = 0.69; NBQX + CPP vs gabazine, p = 0.01; n = 7 cells, 3 animals). These findings indicate that CCK+ interneurons make inhibitory connections onto neighboring L5 pyramidal cells, and that our viral strategy avoids contamination from excitatory contacts due to inadvertent activation of CCK+ pyramidal cells.

**Figure 4:**
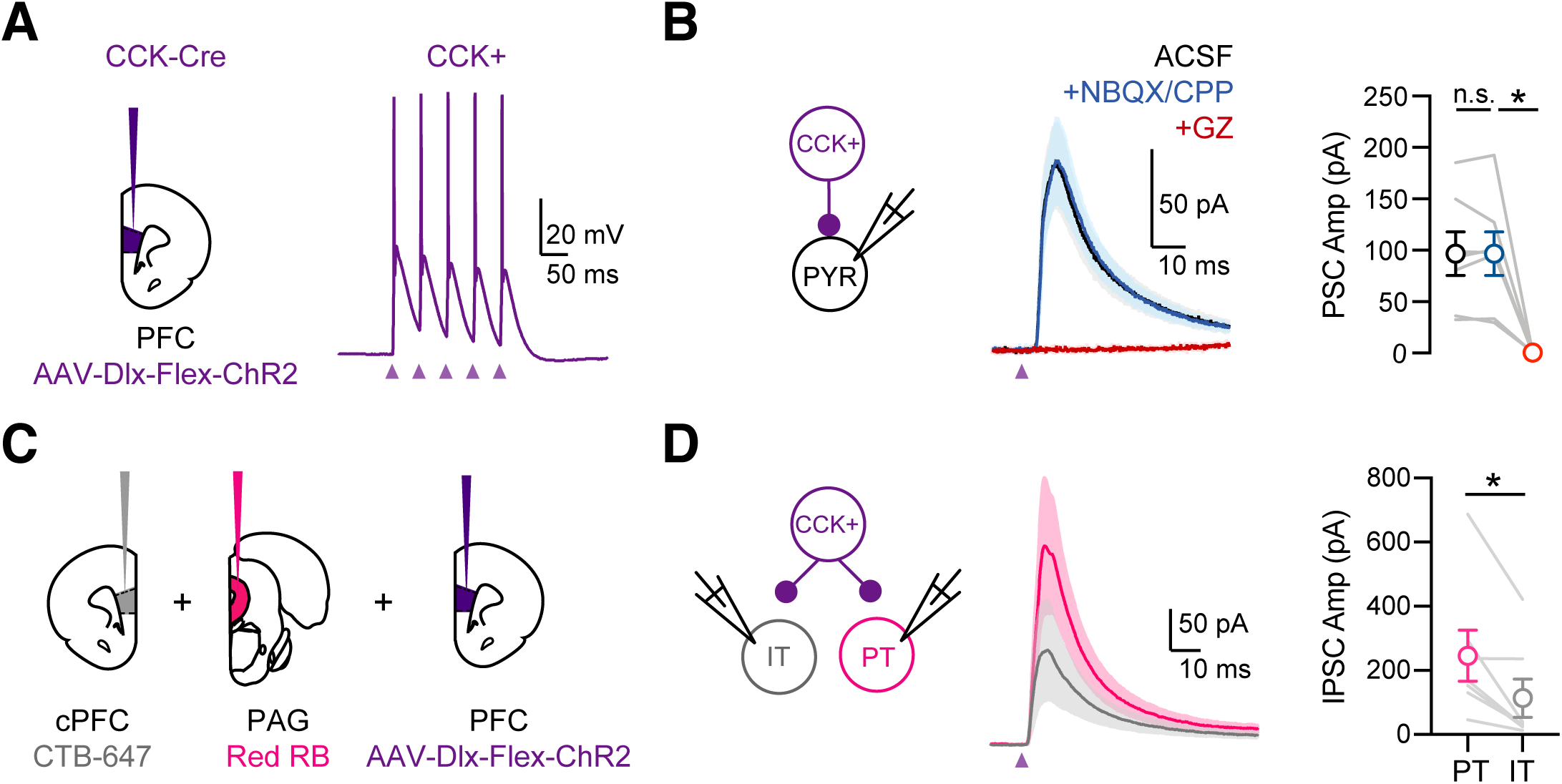
CCK+ interneurons make connections onto pyramidal cells. **A)** *Left*, Injections of AAV-Dlx-Flex-ChR2 into the PFC of CCK-Cre animals. *Right*, Light-evoked firing of a ChR2-expressing CCK+ interneuron. Purple arrows = 5 light pulses at 20 Hz. **B)** *Left,* Average CCK+-evoked IPSCs at L5 pyramidal cells in IL PFC. When recording at −50 mV with a low Cl-internal solution, only outward IPSCs were observed (black). IPSCs were unchanged after wash-in of NBQX + CPP (blue) but abolished by wash-in of gabazine (red). *Right*, Summary of IPSC amplitudes in the different conditions. Purple arrow = light pulse. **C)** Schematic of triple injections, with CTB-647 into cPFC, red retrobeads into PAG, and AAV-Dlx-Flex-ChR2 into PFC. **D)** *Left*, Recording schematic of CCK+ inputs onto IT and PT cells. Middle, CCK-evoked IPSCs at PT and IT cells. *Right*, Summary of IPSC amplitudes at PT and IT cells (7 cells, 4 animals). * p<0.05

Previous studies indicate that inhibitory inputs from PV+ and SOM+ interneurons are strongly biased onto PT cells over nearby IT cells (Anastasiades et al., 2018). To test if similar biases occur for CCK+ interneurons, we labeled PT and IT cells by injecting retrograde tracers into periaqueductal gray (PAG) and cPFC, respectively (**Fig. 4C**). Recording from pairs of pyramidal cells, we found that CCK-evoked IPSCs were larger onto PT cells than IT cells (**Fig. 4D;** IPSC IT = 113 ± 59 pA, PT = 245 ± 79 pA, p = 0.01; n = 7 pairs, 4 animals). Together, these findings indicate our viral strategy can be used to manipulate CCK+ interneurons, and show that these cells make cell-type specific connections, preferentially targeting PT cells in L5 of the IL PFC.

### CB1R-mediated DSI depends on the postsynaptic cell-type

In the hippocampus and the amygdala, CCK+ inputs to pyramidal cells are strongly modulated by endocannabinoids (Lee et al., 2010; Vogel et al., 2016; Wilson and Nicoll, 2001). Postsynaptic depolarization releases endocannabinoids that act on presynaptic CB1 receptors to prevent GABA release, a process known as depolarization-induced suppression of inhibition (DSI) (Wilson and Nicoll, 2001). To study DSI at IT and PT cells, we injected retrogradely transported CTBs in the cPFC and PAG, along with AAV-Dlx-Flex-ChR2 in the PFC of CCK-Cre mice (**Fig. 5A**). We evoked DSI with a standard protocol (Glickfeld and Scanziani, 2006; Wilson and Nicoll, 2001), recording baseline CCK-evoked IPSCs during a train, followed by a 5 s depolarization to +10 mV and recording a DSI test after an initial 1 s delay, followed by a recovery test after another 30 s delay (**Fig. 5B**). For these experiments, it was critical to record IPSCs at −50 mV rather than +10 mV, which allows us to control the timing of endocannabinoid release. In voltage-clamp recordings from retrogradely labeled IT cells, we found that CCK+-evoked IPSCs underwent pronounced DSI that recovered when inputs were stimulated again 30 seconds later (**Fig. 5C & F**; IPSC_1_: before = 158 **±** 34 pA, after = 91 **±** 24 pA, recovery = 149 **±** 33 pA; DSI ratio = 0.57 **±** 0.06, p = 0.004; n = 9 cells, 5 animals). We confirmed the involvement of endocannabinoid signaling by blocking DSI with the CB1R inverse agonist 10 µM AM-251 (**Fig. 5D & F**; IPSC_1_: before = 189 ± 53 pA, after = 168 ± 44 pA, recovery = 185 ± 55; DSI ratio = 0.92 **±** 0.04, p = 0.08; n = 7 cells, 4 animals). We also confirmed this modulation was specific to CCK+ inputs, as PV+ and SOM+-evoked IPSCs at IT cells had minimal DSI (**Fig. 5F & S2**; IPSC_1_ DSI ratio: PV+ = 0.94 **±** 0.02, p = 0.11; n = 7 cells, 3 animals; SOM+ = 0.87 **±** 0.05, p = 0.02; n = 8 cells, 3 animals; PV+ vs CCK+, p = 0.0007; SOM+ vs CCK+, p = 0.008). Surprisingly, we found minimal DSI at PT cells, indicating modulation also strongly depends on postsynaptic cell type (**Fig. 5E & F**; IPSC_1_ before = 208 **±** 40 pA, after = 202 **±** 44 pA, recovery = 205 **±** 40 pA; DSI ratio = 0.95 **±** 0.04, p = 0.22; n = 7 cells, 3 animals). These findings indicate that although CCK+ interneurons broadly contact L5 pyramidal cells, the connections undergo prominent DSI only at IT cells and not nearby PT cells.

**Figure 5:**
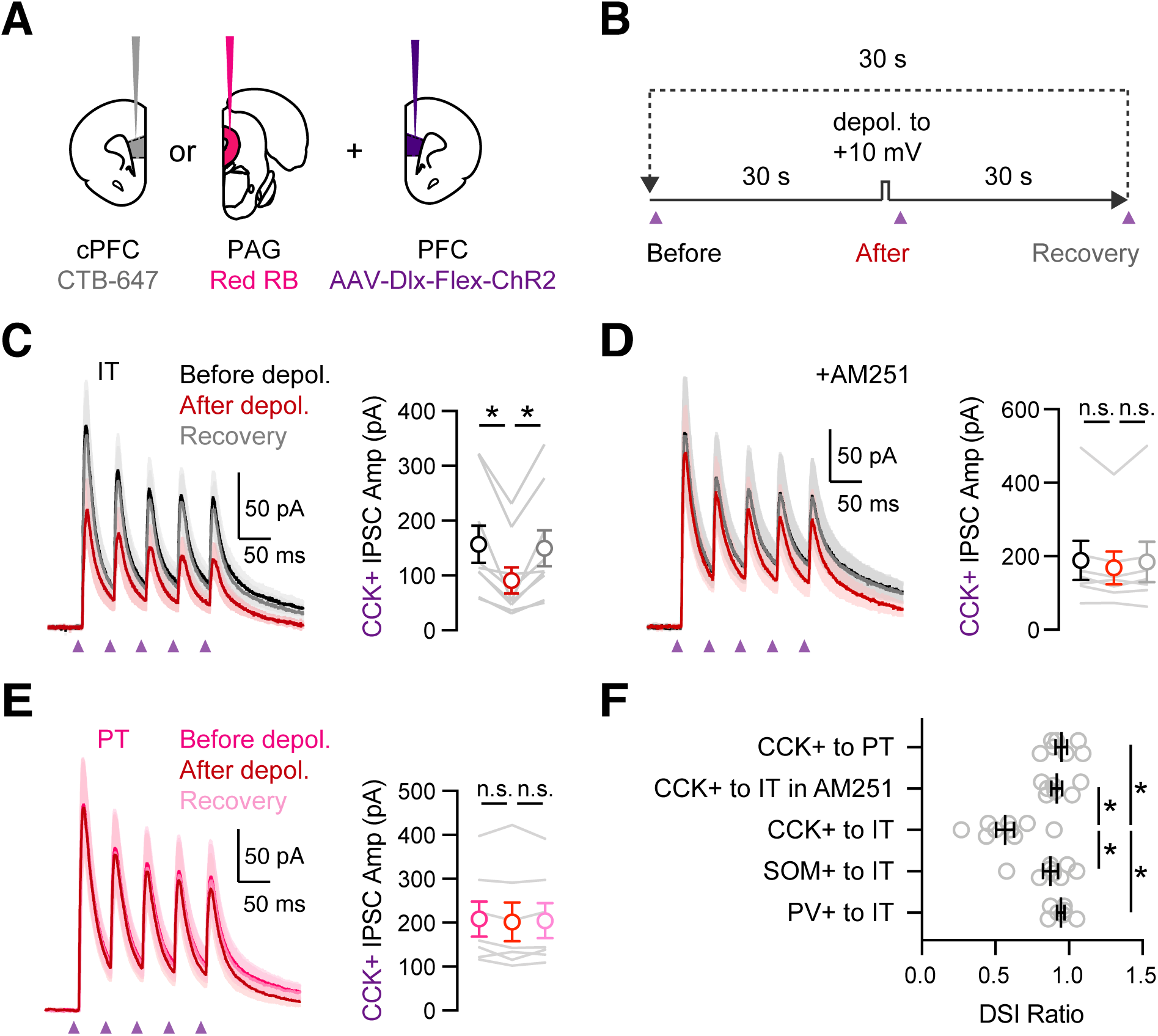
Prominent DSI of CCK+ inputs onto IT cells. **A)** Injection schematic of CTB-647 into cPFC and red retrobeads (RB) into PAG, along with AAV-Dlx-Flex-ChR2 into PFC of CCK-Cre mice. **B)** Experimental procedure for depolarization-induced suppression of inhibition (DSI), with 30 s baseline, followed by 5 s depolarization to +10 mV, and 30 s recovery, repeated every 30 s. **C)** *Left*, Average CCK+-evoked IPSCs at IT cells before (black), after (red) and recovery from (gray) the brief depolarization. Purple arrows = 5 light pulses at 20 Hz. *Right*, Summary of amplitudes of the first IPSC, showing robust DSI (n = 9 cells, 5 animals). **D)** Similar to (C) in the presence of 10 µM AM-251, which abolished DSI (n = 7 cells, 4 animals). **E)** Similar to (C) for CCK+-evoked IPSCs at PT cells, showing no DSI (n = 7 cells, 3 animals). **F)** Summary of DSI ratios (IPSC after depolarization / IPSC before depolarization) across experiments in (C – E). * p<0.05

### Endocannabinoid modulation depends on postsynaptic cell type

The dependence of DSI on postsynaptic cell-type could have a presynaptic origin if CB1Rs are expressed on a subset of CCK+ connections. To test this possibility, we next used immunocytochemistry to examine CB1R+ puncta at the cell bodies of IT and PT cells. To label these projection neurons, we injected retrogradely transported AAVrg-TdTomato into the cPFC and AAVrg-GFP into the PAG, respectively (**Fig. 6A**). While we observed CB1R+ puncta in L5 around the cell bodies of both cell types (**Fig. 6A**), their density was greater at IT cells (**Fig. 6B;** IT = 9.3 ± 0.5 puncta / cell, PT = 5.7 ± 0.5 puncta / cell, p < 0.0001; n = 4 animals, 247 IT cells, 207 PT cells). These results indicate that presynaptic CB1Rs are more prominent at perisomatic connections onto IT cells, suggesting why these inputs are more strongly affected by DSI.

**Figure 6:**
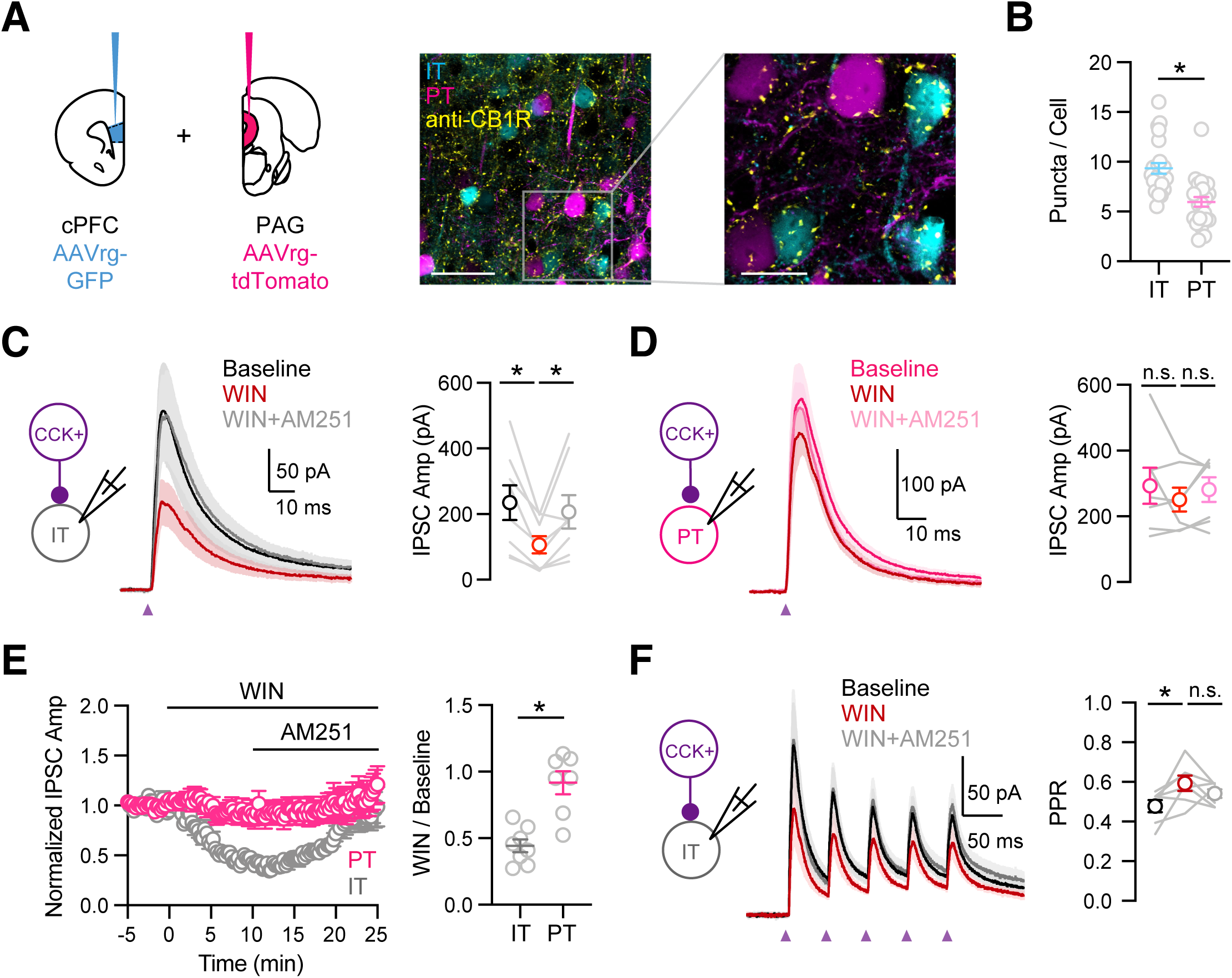
Cell-type specific modulation by CB1 receptors. **A)** *Left*, Injection schematic of AAVrg-GFP into PFC and AAVrg-tdTomato into PAG. *Middle*, Confocal image of IT cells (cyan), PT cells (purple) and CB1 receptors (yellow). Scale bar = 50 µm. *Right*, Magnification of region on left. Scale bar = 20 µm. **B)** Quantification of CB1R puncta in IT and PT cells, each dot represents the average puncta number per cell in a slice (n = 247 IT cells, 207 PT cells, 4 animals). **C)** *Left,* Schematic of recordings from IT cells in L5 of IL PFC. *Middle,* Average CCK+-evoked IPSCs at IT cells at baseline (black), 10 min after wash-in of 1 µM WIN 55,212-2 (red), and 15 minutes after additional wash-in of 10 µM AM-251 (gray). Purple arrow = light stimulation. *Right*, Summary of IPSC amplitudes (n = 8 cells, 5 animals). **D)** Similar to (C) for CCK+-evoked IPSCs at PT cells, showing lack of modulation by CB1R (n = 7 cells, 4 animals). **E)** *Left,* Summary of time course of modulation at IT and PT cells, with IPSC amplitudes normalized to the average response during the first 5 minutes. *Right,* Summary of normalized IPSC amplitudes after WIN wash-in. **F)** Similar to (C) for trains of CCK+ inputs onto IT cells (5 pulses at 20 Hz), showing small increase in PPR after wash-in of 1 µM WIN 55,212-2 (n = 7 cells, 4 animals). * p<0.05

If the cell-type specificity of DSI depends on presynaptic factors, we would expect to observe equivalent differences for pharmacologically evoked endocannabinoid modulation. We further examined the mechanism of CB1R modulation using wash-in of the agonist WIN 55,212-1 (WIN, 1 µM) followed by the inverse agonist AM-251(10 µM), which act directly at presynaptic CB1Rs located on CCK+ axon terminals. In voltage-clamp recordings from L5 IT cells, we found that WIN reduced CCK+-evoked IPSCs, which was reversed by AM-251 (**Fig. 6C**; baseline = 234 ± 53 pA, WIN = 106 ± 26 pA, AM-251 = 206 ± 50 pA; baseline vs WIN, p = 0.008; WIN vs AM-251, p = 0.016; n = 8 cells, 5 animals). In contrast, in recordings from PT cells, we found that neither WIN nor AM-251 had any effect on CCK+-evoked IPSCs, consistent with their lack of DSI (**Fig. 6D**; baseline = 292 ± 55 pA, WIN = 250 ± 36 pA, AM-251 = 281 ± 37 pA; baseline vs WIN, p = 0.81; WIN vs AM-251, p = 0.22; n = 7 cells, 4 animals). Overall, activation of CB1 receptors strongly reduced CCK+-evoked IPSCs only at IT cells but not neighboring PT cells (**Fig. 6E**; WIN / baseline: IT = 0.44 ± 0.05, PT = 0.92 ± 0.09, p = 0.001). Interestingly, during trains of CCK+ stimulation, wash-in of WIN also increased the paired-pulse ratio (PPR) at IT cells, suggesting presynaptic modulation by CB1Rs (**Fig. 6F**; IPSC_2_ / IPSC_1_: baseline = 0.48 ± 0.03, WIN = 0.59 ± 0.03, AM-251 = 0.54 ± 0.02; baseline vs WIN, p = 0.03; WIN vs AM-251, p = 0.22; n = 7 cells, 4 animals). These findings demonstrate that differences in presynaptic endocannabinoid signaling can account for the differential presence of DSI at CCK+ connections onto IT and PT cells.

### vHPC-evoked feed-forward inhibition is modulated by endocannabinoids

Together, our results suggest that vHPC inputs strongly engage CCK+ interneurons that in turn robustly inhibit IT cells. Based on the presence of CB1R modulation and DSI at IT cells, we hypothesized that vHPC-evoked inhibition should also undergo target-specific DSI. In voltage-clamp recordings from IT cells, we found that repetitive activation of vHPC inputs evoked EPSCs and IPSCs at −50mV (**Fig. 7A & B**). Depolarization of IT cells (5 s to +10 mV) reduced vHPC-evoked IPSCs but not EPSCs at −50 mV (**Fig. 7B & E**; before, after, recovery: IPSC_1_ = 102 ± 14 pA, 51 ± 10 pA, 88 ± 10 pA; EPSC_1_ = 248 ± 54 pA, 233 ± 48 pA, 241 ± 52 pA; DSI ratio = 0.47 ± 0.05, p = 0.02; DSE ratio = 0.99 ± 0.06, p = 0.30; n = 7 cells, 4 animals). These findings indicate that there is no DSE at the vHPC to PFC connection, whereas there is prominent DSI onto IT cells. Importantly, application of AM-251 minimized DSI, indicating that it is mediated by endocannabinoids activating CB1Rs (**Fig. 7C & E**; IPSC_1_: before = 136 **±** 31 pA, after = 116 **±** 29 pA, recovery = 149 **±** 35 pA; DSI ratio = 0.84 **±** 0.03, p = 0.02; n = 7 cells, 3 animals; DSI in ACSF vs AM-251, p = 0.0006). Moreover, depolarization of PT cells had minimal effect on either vHPC-evoked IPSCs or EPSCs, confirming no DSI or DSE onto this postsynaptic cell type (**Fig. 7D & E**; IPSC_1_ before = 129 **±** 32 pA, after = 114 **±** 27 pA, recovery = 134 **±** 33 pA; DSI ratio = 0.91 ± 0.05, p = 0.10; DSE ratio = 0.89 ± 0.04, p = 0.01; n = 10 cells, 4 animals). These findings show that DSI of vHPC-evoked inhibition only occurs at IT cells, suggesting a key role for CCK+ interneurons in the communication between vHPC and PFC.

**Figure 7:**
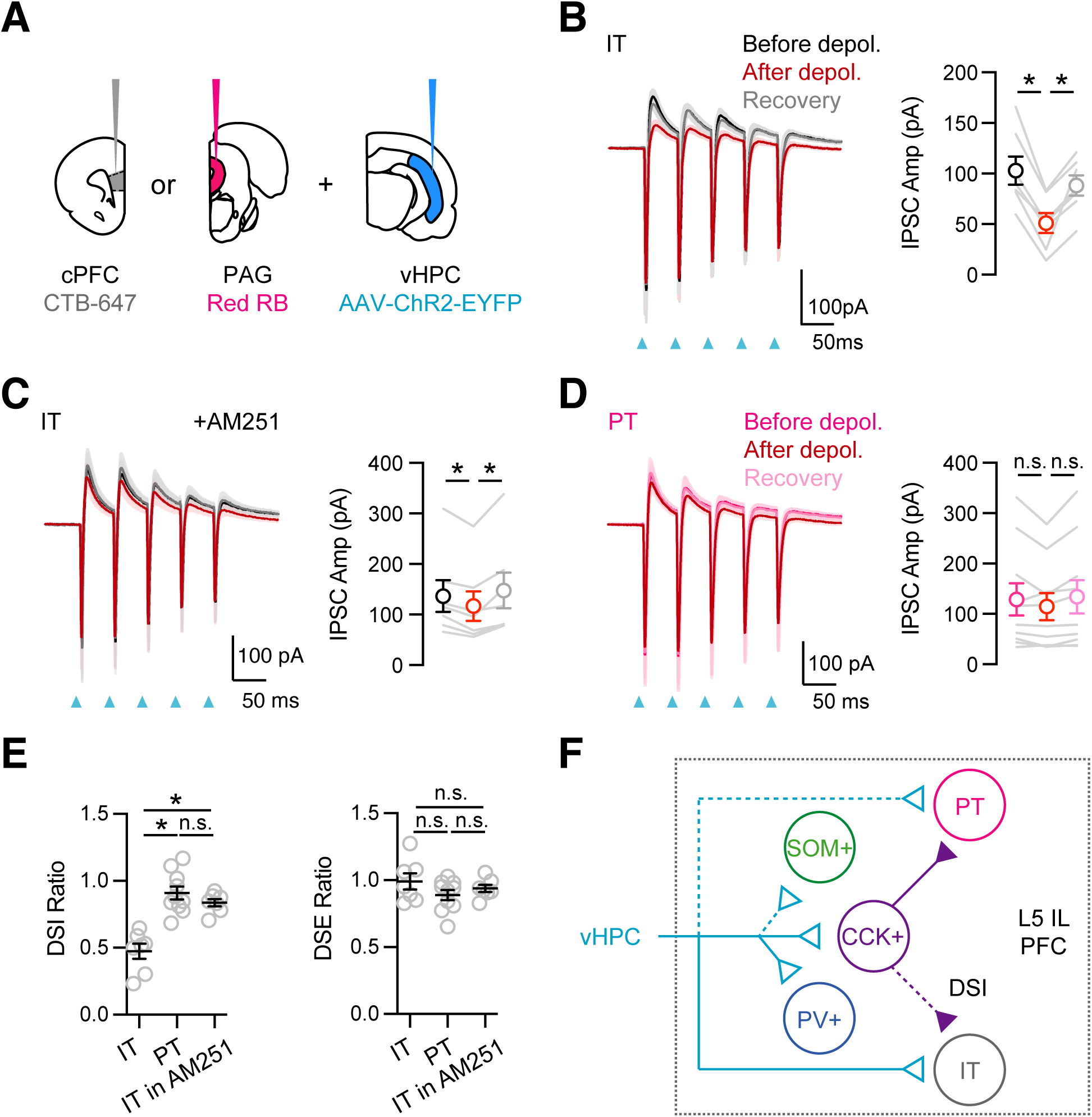
vHPC-evoked feed-forward inhibition at IT cells undergoes DSI. **A)** Injection schematic, showing CTB-647 in cPFC, red retrobeads (RB) in PAG, along with AAV-ChR2-EYFP in vHPC. **B)** *Left*, Average vHPC-evoked EPSCs and IPSCs at IT cells in L5 IL PFC, before (black), immediately after depolarization (red), and after recovery (gray) (same paradigm as Figure 5). Blue arrows = light stimulation. *Right*, Summary of amplitudes of first vHPC-evoked IPSCs (n = 7 cells, 4 animals). **C)** Similar to (B) in the presence of 10 µM AM-251, which reduces DSI (n = 7 cells, 3 animals). **D)** Similar to (B) for PT cells, showing no DSI (n = 10 cells, 4 cells). **E)** Summary of DSI and DSE ratios (amplitude ratios of IPSCs or EPSCs after / before the depolarizations) across the different experiments. **F)** Summary schematic for vHPC-evoked feed-forward inhibition in IL PFC. vHPC inputs directly contact IT over PT cells to evoke EPSCs. vHPC inputs also engage multiple interneurons to evoke local inhibition. Inhibition mediated by CCK+ interneurons displays robust endocannabinoid-mediated DSI, but only at IT cells, and not neighboring PT cells. * p < 0.05

## DISCUSSION

We have explored several new features about the organization and modulation of connections from vHPC to PFC (**Fig. 7F**). First, we found vHPC contacts and strongly activates CB1R-expressing CCK+ interneurons in L5 of IL PFC. Second, we showed that CCK+ interneurons contact nearby pyramidal cells, suggesting they participate in feed-forward inhibition. Third, we found that CCK+ connections undergo CB1R-mediated modulation and DSI, which is selective for IT and not PT cells. Fourth, endocannabinoids also modulate vHPC-evoked inhibition, which also undergoes DSI selectively at IT cells. Together, our results reveal a central role for CCK+ interneurons and endocannabinoid modulation in communication between vHPC and PFC.

The PFC possesses a rich variety of GABAergic interneurons, which are known to have unique roles in goal-directed behaviors (Abbas et al., 2018; Courtin et al., 2014; Kepecs and Fishell, 2014; Kvitsiani et al., 2013). Interestingly, the PFC has fewer PV+ interneurons and more CCK+ interneurons compared to other cortices (Kim et al., 2017; Whissell et al., 2015). Our results indicate that CCK+ interneurons are abundant in L5 of IL PFC, which we previously showed receives the strongest connections from vHPC (Liu and Carter, 2018). We found these cells are distinct from PV+ and SOM+ interneurons, with different morphological and physiological properties. We also found that they are enriched in presynaptic CB1Rs, as in other parts of cortex, hippocampus and amygdala (Armstrong and Soltesz, 2012; Bodor et al., 2005; Vogel et al., 2016). The enrichment of CCK+ interneurons across limbic areas is consistent with a role in emotional behaviors (Freund and Katona, 2007; Klausberger et al., 2005). In the future, it will be interesting to characterize the properties of CCK+ interneurons in other layers and subregions of PFC.

One of our key results is that vHPC inputs densely contact and strongly activate CCK+ interneurons in L5 of IL PFC. vHPC inputs are strong and depressing onto CCK+ interneurons, in contrast to the facilitating inputs onto IT cells (Liu & Carter, 2018). Importantly, vHPC-evoked firing of CCK+ interneurons occurs without activation of IT cells, indicating polysynaptic recurrent activity is not required for activation of CCK+ interneurons, which are therefore likely to mediate feed-forward inhibition. With higher intensity of vHPC stimulation, CCK+ interneurons may also be activated by local inputs, as observed in the hippocampus (Glickfeld and Scanziani, 2006), and similar to SOM+ interneurons in the cortex (Silberberg and Markram, 2007). Indeed, in the hippocampus, CCK+ interneurons have been shown to participate in both feed-forward and feed-back networks (Basu et al., 2013; Glickfeld and Scanziani, 2006). Our findings show that CCK+ interneurons have an important and underappreciated role in hippocampal evoked feed-forward inhibition, and it will be important to assess whether this generalizes to other inputs to the PFC.

While we focused on CCK+ interneurons, we also confirmed that vHPC inputs engage other interneurons in L5 of IL PFC. Activation of PV+ interneurons is strong, consistent with their role in feed-forward inhibition in PFC and elsewhere (Anastasiades et al., 2018; Cruikshank et al., 2007; Delevich et al., 2015; Gabernet et al., 2005; McGarry and Carter, 2016). Previous results have indicated that vHPC engages PV+ interneurons in superficial layers of the medial PFC (Marek et al., 2018). In contrast, we find particularly strong connections in L5, where our previous study shows vHPC inputs are most dense (Liu and Carter, 2018). SOM+ interneuron activation is weak, but increases with repetitive activity, similar to BLA inputs to superficial PFC (McGarry and Carter, 2016). The engagement of SOM+ interneurons is consistent with a role in oscillations linking the vHPC and PFC and involvement in working memory (Abbas et al., 2018). Interestingly, the activation of SOM+ interneurons by long-range inputs also occurs in granular sensory cortex, where it also builds during stimulus trains, suggesting this is a general property (Tan et al., 2008).

We found that CCK+ interneurons make strong inhibitory connections onto neighboring pyramidal cells in L5 of the IL PFC. Importantly, these connections are much stronger onto PT cells, similar to our previous findings for PV+ and SOM+ interneurons (Anastasiades et al., 2018). Biased inhibition is thus a general property of inhibitory connections in the PFC, although strength also depends on intrinsic properties (Anastasiades et al., 2018). In the hippocampus, CCK+ interneurons also make unique connections onto different pyramidal cell populations, although these are typically defined by sublayer (Valero et al., 2015). Studying the output of CCK+ interneurons was enabled by using Cre-dependent viruses with Dlx enhancers (Dimidschstein et al., 2016). These tools allowed us to avoid simultaneous activation of pyramidal cells, which also express CCK (Taniguchi et al., 2011). In the future, this approach can be used to study how CCK+ interneurons contact other cells in the local network. For example, CCK+ interneurons may also target other classes of interneurons, as observed in the hippocampus (Daw et al., 2009).

Another key finding of our work was that endocannabinoid modulation of CCK+ connections depends on the postsynaptic target. We observe robust DSI at IT cells, but minimal DSI at nearby PT cells, despite the latter receiving stronger connections. This cell-type specific DSI has rarely been examined in the cortex, as most studies do not distinguish projection target. However, a recent study also showed differential DSI at CCK+ connections onto projection neurons in the amygdala (Vogel et al., 2016). The dependence of DSI on the postsynaptic target should be taken into account in future studies of CCK+-mediated inhibition. Importantly, our experiments suggested no equivalent DSI for either PV+ or SOM+ connections onto IT cells. These results suggest CCK+ connections are a primary target of endocannabinoid modulation, in agreement with previous studies in the cortex, hippocampus and other brain regions (Bodor et al., 2005; Glickfeld et al., 2008; Gonchar et al., 2007; Katona et al., 1999; Kawaguchi and Kondo, 2002).

In principle, the selectivity of DSI at CCK+ connections onto IT cells could reflect differences in postsynaptic release or presynaptic detection of endocannabinoids. Our results are consistent with the latter explanation. First, our immunocytochemistry shows more CB1R puncta around IT cells, suggesting these receptors are selectively localized. This result was particularly surprising because we also found CCK+ interneurons make stronger connections onto PT cells. Second, direct activation of CB1Rs with WIN reduces CCK-evoked IPSCs only at IT cells, with no effect at neighboring PT cells. This experiment bypasses postsynaptic release of endocannabinoids, suggesting a presynaptic mechanism can account for specificity of DSI. The increase in PPR is also consistent with presynaptic modulation by CB1R, suggesting reduced release probability (Wilson et al., 2001). This work underscores that endocannabinoid modulation is cell-type specific, and it will be important to examine if similar specificity occurs in other regions of the PFC.

The ability of vHPC inputs to engage CCK+ interneurons, which in turn contact pyramidal cells, implicates a key role in feed-forward inhibition. Consistent with this idea, we observed prominent CB1R-mediated DSI of vHPC-evoked inhibition only at IT cells and not neighboring PT cells. Because there is no change in excitation, this will selectively increase the excitation / inhibition (E / I) ratio at IT cells. This could allow the vHPC to more effectively activate IT cells compared to neighboring PT cells, perhaps elevating local processing within the PFC. Altered E / I balance could also potentially shift PFC output to other intratelencephalic targets throughout the brain, including targets in the cortex, striatum, amygdala and claustrum (Anastasiades et al., 2019). Together, our findings suggest that both CCK+ interneurons and endocannabinoid modulation play a fundamental role in vHPC to PFC communication. In the future, it will be important to assess the contribution of this circuit to emotional control and disorders, including threat learning and anxiety disorders (Peters et al., 2010; Sierra-Mercado et al., 2011; Sotres-Bayon et al., 2012).

## Acknowledgements

We thank the Carter lab for helpful discussions and comments on the manuscript. This work was supported by NIH R01 MH085974 (AGC). The authors have no financial conflicts of interest.

---

**Figure S1:**
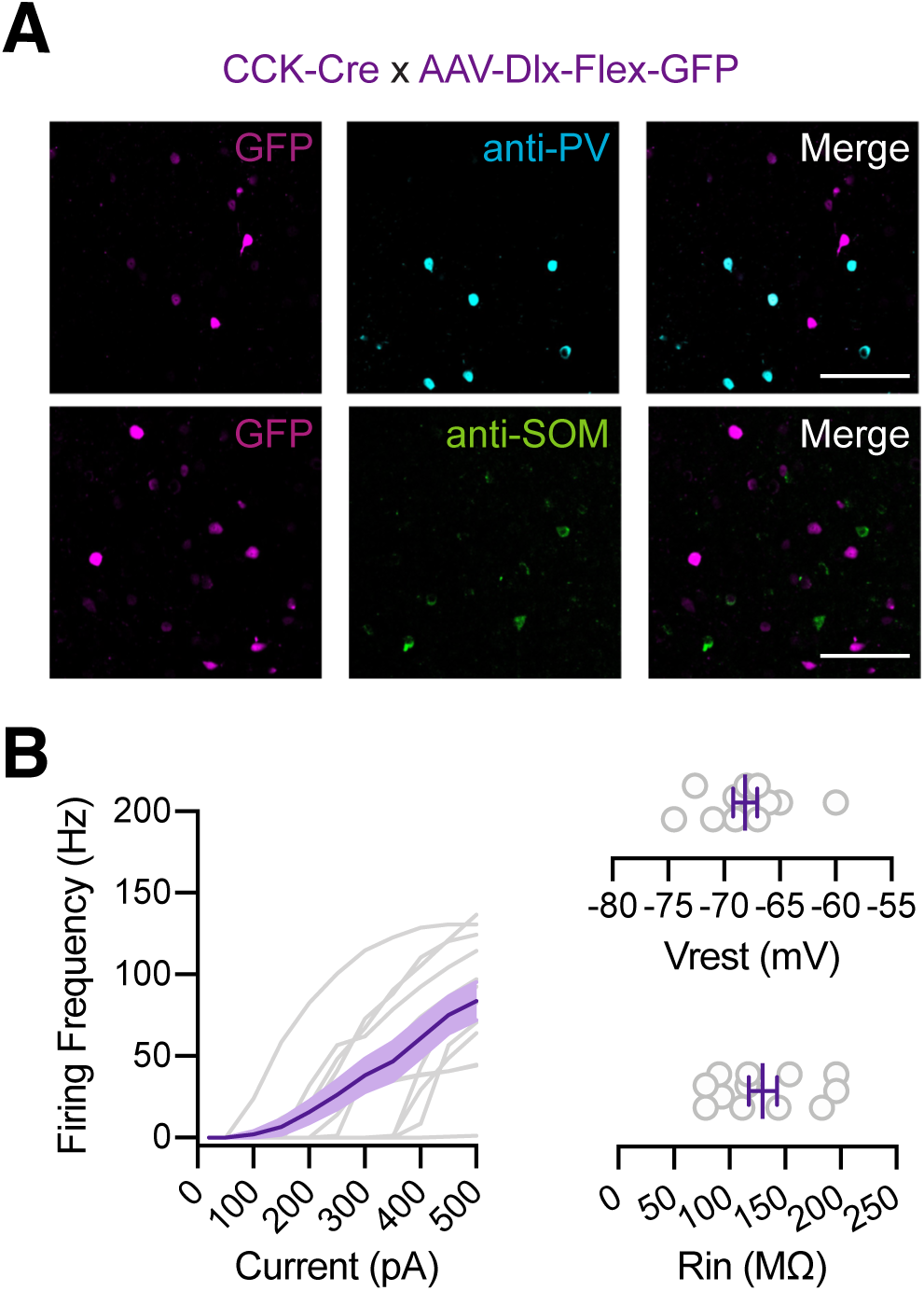
Properties of CCK+ interneurons in L5 IL PFC. **A)** Co-labeling of GFP-expressing CCK+ interneurons (purple) with PV (blue) and SOM (green). Immunohistochemistry of GFP and either PV (top) or SOM (bottom) in PFC slices from CCK-Cre animals injected with AAV-Dlx-Flex-GFP. From left to right: GFP; PV or SOM; merged. Scale bar = 100 µm. **B)** *Left*, Firing frequency (F) vs. current (I) curve for CCK+ cells. *Right,* Summary of membrane resting potential (Vrest) and input resistance (Rin) of CCK+ interneurons (n = 12 cells, 4 animals).

**Figure S2:**
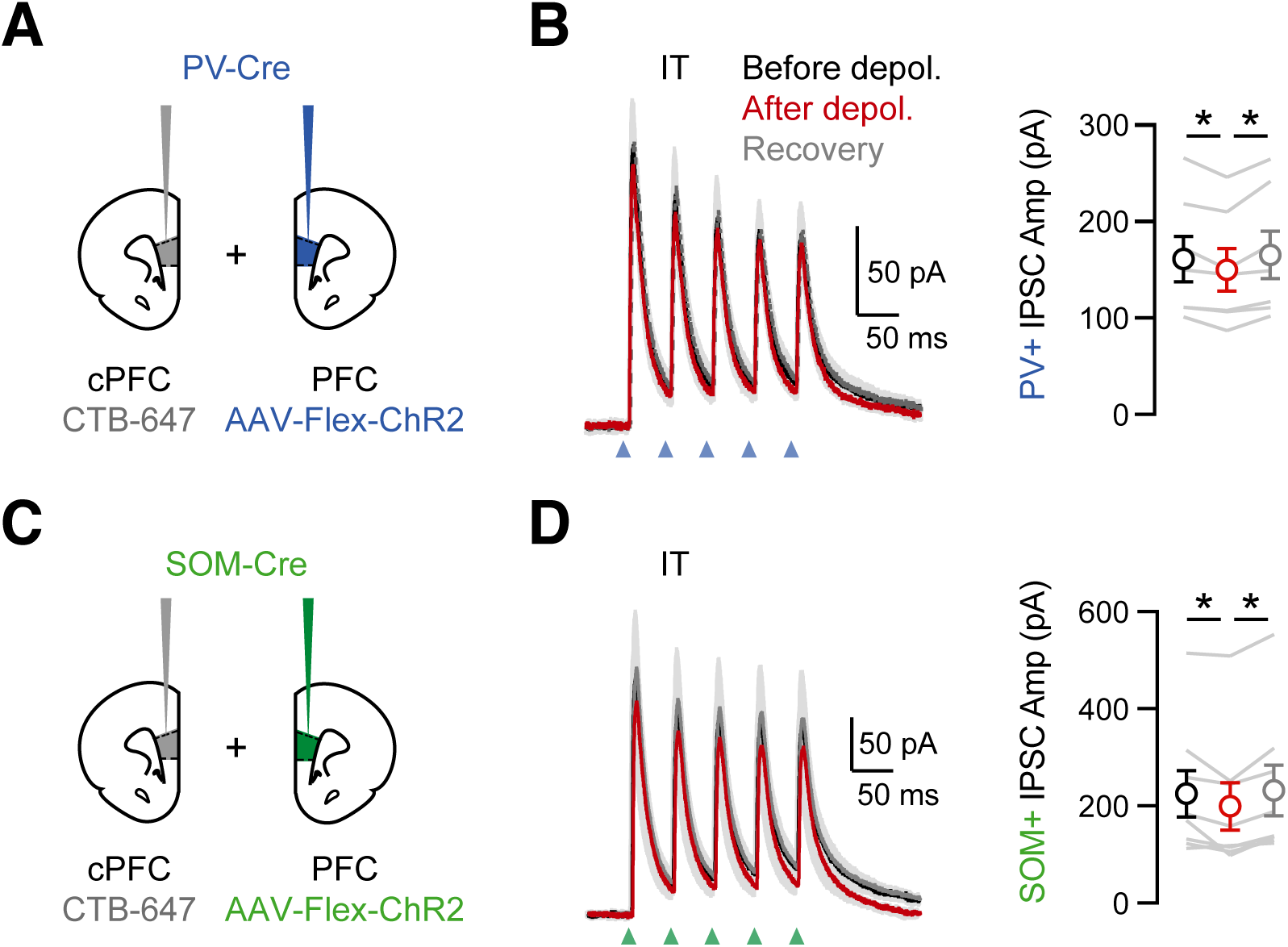
Minimal DSI at PV+ or SOM+ interneuron connections onto IT cells. **A)** Injection schematics of CTB-647 into cPFC and AAV-Flex-ChR2 in PFC of PV-Cre mice. **B)** *Left*, Average PV+ inputs onto IT cells, evoked by 5 pulses at 20 Hz, before (black), immediately after the brief depolarization (red) and after recovery (gray). Blue arrow = light stimulation. *Right*, Amplitudes of the first IPSCs (n = 7 cells, 3 animals). **C – D)** Similar to (A – B) for SOM+ inputs onto IT cells (n = 8 cells, 3 animals).

